# PPAR-γ is a promising therapeutic target for memory deficits induced by early alcohol exposure

**DOI:** 10.1101/2023.01.20.524912

**Authors:** Alba Garcia-Baos, Antoni Pastor, Ines Gallego-Landin, Rafael de la Torre, Ferran Sanz, Olga Valverde

**Affiliations:** Neurobiology of Behavior Research Group (GReNeC-NeuroBio), Department of Medicine and Life Sciences, Universitat Pompeu Fabra, Barcelona, Spain; Neuroscience Research Program, IMIM-Hospital del Mar Research Institute, Barcelona, Spain; Research Program on Biomedical Informatics (GRIB), IMIM-Universitat Pompeu Fabra, Barcelona, Spain

**Keywords:** fetal alcohol spectrum disorder, memory deficits, expanded endocannabinoid system, peroxisome proliferator-activated receptor type gamma, astrocytes, hippocampus

## Abstract

**Background:** Patients diagnosed with fetal alcohol spectrum disorder (FASD) show persistent cognitive disabilities, including memory deficits. However, the neurobiological substrates of these deficits remain unclear. Here, we studied the participation of the expanded endocannabinoid system (ECS), which is known to be affected by alcohol in other life periods, and it is involved in memory impairments of neurodevelopmental disorders.

**Methods:** C57BL/6 female mice were exposed to a time-limited access to either water or alcohol to model prenatal and lactation alcohol exposure (PLAE). The expanded ECS was analyzed in the prefrontal cortex and the hippocampus of the offspring at post-partum day (PD) 25 and 70. Then, memory performance was tested after the repeated administration (from PD25 to PD34) of: i) URB597, to increase N-acylethanolamines (NAEs), and GW9662, a peroxisome proliferator-activated receptor gamma (PPAR-γ) antagonist; ii) pioglitazone, a PPAR-γ agonist. Finally, we used a viral approach to upregulate astrocytic PPAR-γ in the hippocampus to restore memory deficits.

**Results:** We report that PLAE causes a hippocampal reduction of NAEs and PPAR-γ at PD25. Moreover, URB597 suppresses PLAE-induced memory deficits through PPAR-γ, since its effects are prevented by GW9662. Direct PPAR-γ activation, using pioglitazone, also ameliorates memory impairments. Lastly, we demonstrate that the upregulation of PPAR-γ in hippocampal astrocytes is sufficient to rescue PLAE-induced memory deficits.

**Conclusion:** Our data reveal a bidirectional link between memory deficits and expanded ECS alterations in the context of early alcohol exposure. Furthermore, we proved that PPAR-γ in hippocampal astrocytes represents a specific therapeutic target for memory deficits in FASD.

## Introduction

Perinatal alcohol exposure produces a wide range of physical and behavioral disabilities comprised under the term fetal alcohol spectrum disorder (FASD). Cognitive impairments are the most prevalent and disabling symptoms although clinical manifestations significantly differ among patients (1,2). Perinatal alcohol exposure disrupts major developmental processes, like neuro- and gliogenesis, migration, proliferation, synaptogenesis and myelination (3–6), consequently impairing synaptic plasticity, circuit formation, learning and memory (7). These molecular and behavioral alterations are physiologically irreversible and persistent (8–11). Therefore, an early intervention during critical periods of high brain plasticity could facilitate the resolution of the disrupted processes on time and ameliorate the prognosis.

The expanded endocannabinoid system (ECS) is a highly complex network that includes classic endocannabinoids and receptors, but also other related molecules and metabolic pathways (12). Given that it is widely expressed over the central nervous system since very early developmental stages, this modulatory system participates in crucial maturation processes (13,14). Its main elements include endocannabinoids and other related lipid mediators, such as N-acylethanolamines (NAEs) and 2-acylglycerols; the enzymatic machinery that synthetize and degrade said endocannabinoids and other related lipid mediators; and the receptors that mediate their biological activities (12). Besides canonical cannabinoid receptor (CB)1 and CB2, other receptors like peroxisome proliferator-activated receptor type gamma (PPAR-γ) are involved in these biological effects (12,15,16). PPAR-γ is a nuclear receptor that, when activated, migrates to the nucleus to modulate the gene expression (17). Importantly, it is involved in processes that are key for brain maturation and cognition (18,19). In fact, hippocampal alterations of PPAR-γ during neurodevelopment have been consistently associated to cognitive impairments (19–21). In addition, NAEs and exogenous cannabinoids exert neuroprotective effects via PPAR-γ activation in the context of cognitive dysfunction (15,22). Given its lipidic nature, the expanded ECS is remarkably compromised by alcohol exposure (23,24). Indeed, perinatal alcohol exposure in mice produces changes in AEA levels and CB1 expression at least in neocortical and hippocampal regions (25). Even though studies of perinatal alcohol exposure on non-canonical molecules of the expanded ECS are limited, some evidence does suggest that this complex system might provide new therapeutic opportunities for FASD. On the one hand, the administration of cannabidiol, which acts on multiple targets of the expanded ECS including NAEs and PPAR-γ, ameliorates cognitive deficits in a FASD-like mouse model (9,26) and in FASD patients (27,28). On the other hand, resveratrol, a natural PPAR-γ agonist, attenuated cognitive impairments in a preclinical model of FASD (29). Likewise, *in vitro* studies show that the selective PPAR-γ agonist pioglitazone induced neuroprotective effects through anti-inflammatory mechanisms in a FASD-like mouse model (10,30). However, the exact mechanisms underlying its protective effects on memory remain unclear.

The hippocampus (HPC) is a primary region involved in memory function, and hippocampal cells are particularly vulnerable to early alcohol exposure (31). Notably, astrocytic cells play an essential role in brain development under physiological conditions (32,33), contributing to the maintenance of cerebral homeostasis and the protection of neurons from environmental insults (34,35). Astrocytes and astrocytic-mediated neurodevelopmental processes in the HPC are damaged by alcohol exposure, as reported in *in vitro* studies of astrocytic and neuronal cultures from embryonic rats (36,37). Moreover, hippocampal synaptic plasticity, shaped by astrocytes (38), and required for memory encoding and recall, is dramatically impaired in experimental models of FASD (39,40). Thus, given the peri-synaptic location of astrocytic cells, their immunoreactive capacities, and the fact that astrogenesis occurs after neurogenesis and concurrently with neuronal synaptogenesis (41,42), astrocytes are interesting candidates for the treatment of neurodevelopmental disorders. For this, further understanding of molecular targets in astrocytes that ameliorate core symptoms of FASD is essential. Knowing that early alcohol exposure disrupts brain lipid homeostasis (43,44), and that astrocytes express both NAEs (45) and PPAR-γ (46,47), we propose that targeting the expanded ECS, and more specifically, PPAR-γ in hippocampal astrocytes could be a promising approach to further comprehend how early alcohol exposure persistently alters memory function.

Consequently, we investigated the relationship between alterations in the expanded ECS and memory impairments induced by a mouse model of prenatal and lactation alcohol exposure (PLAE). In the present work, we show that PPAR-γ activation, either by NAEs or pioglitazone, rescues memory deficits in PLAE mice. In addition, we provide evidence that PPAR-γ in hippocampal astrocytes is a specific target with therapeutic potential for memory deficits caused by perinatal alcohol exposure.

## Methods and Materials

### 1. Experimental subjects

Male and female C57BL/6 were purchased from Envigo (Barcelona, Spain) and transported to the animal facility (UBIOMEX, PRBB) to be used as breeders. Animals were housed in a room with controlled temperature (21 ± 1 °C), humidity (55 ± 10%) and lighting (lights on between 7:30 p.m. and 7:30 a.m.). Animals were 12 weeks old when the breeding began. For this, each male was housed with 2 females. After successful mating, pregnant females were individualized and observed daily for parturition. For each litter, the date of birth was designated as post-partum day (PD) 0. Pups remained with their mothers until weaned at PD21. After weaning, male and female offspring were housed separately. Food and water were available *ad libitum*. All animal care and experimental procedures were conducted in accordance with the European Union Directive 2010/63/EU regulating animal research and were approved by the local Animal Ethics Committee (CEEA-PRBB).

### 2. General experimental design

The Drinking in the Dark (DID) procedure was conducted as previously reported (3,9) to model a prenatal and lactation alcohol exposure under a binge-like drinking pattern. After the weaning, we performed four different experiments in PLAE and control (water-exposed) offspring: i) analyses of molecules of the expanded ECS in the HPC and PFC at PD25 and PD70. ii) Based on these molecular results, we performed two pharmacological approaches, in which treatments were administered from PD25 to PD34. In one group of mice, the inhibitor of fatty acid amide hydrolase enzyme (FAAH) URB597 was co-administered with GW9662, a PPAR-γ antagonist, to elucidate if increasing the levels of NAEs had any effects on PLAE-induced memory impairments, and if so, whether these effects were mediated by PPAR-γ. In a second group of mice, pioglitazone, a PPAR-γ agonist, was administered to explore whether direct PPAR-γ activation is sufficient to ameliorate memory deficits caused by PLAE. All treatments were followed by a battery of memory tests starting at PD60 (see timelines in Fig3.A,E). iii) Anatomical description by immunohistochemistry to explore whether PLAE decreases hippocampal PPAR-γ in astrocytes and/or neurons. iv) Based on the anatomical data, we used a viral approach to specifically overexpress PPAR-γ in hippocampal astrocytes, followed by the battery of memory test starting at PD60 (see timeline in Fig5.C).

Animals were always randomly assigned to the experimental groups and treatments to avoid litter effects. Male and female offspring were used as a whole population since no significant differences between sexes were observed in preliminary data from our team. Nonetheless, all the experimental groups contained a balanced number of male and female to avoid bias.

All procedures are detailed in Supplemental Materials.

### 3. Drugs

Ethyl alcohol (Merck Chemicals, Darmstadt, Germany) was diluted in tap water to obtain a 20% (v/v) alcoholic solution. URB597, GW9662 and Pioglitazone hydrochloride (Merck Life Science S.L.U., Madrid, Spain) were diluted in 2% Tween-80 and saline solution (0.9% NaCl). The doses of URB597 (0.3 mg/kg, i.p.) (48–51) and pioglitazone (PIO, 10 mg/kg, i.p.) (52–54) were chosen based on previous studies proving beneficial effects on other pathological animal models. The dose of GW9662 (1 mg/kg, i.p.) was selected because innocuous effects have been shown in other pharmacological studies (55–57).

### 4. Statistics

Student’s unpaired t-tests were applied to evaluate differences between water-exposed and PLAE mice in: i) quantification of endocannabinoids and other NAEs ii) gene expression of receptors and enzymes related to the expanded ECS, iii) immunohistochemical analyses of PPAR-γ, PPAR-γ/GFAP, and PPAR-γ/NeuN positive cells. Two-way analyses of variance (ANOVA) were used to assess the effects of the main factors and their interactions on the reference memory, NOR and NOL tests. We applied these analyses for the behavioral experiments assessing the effects of PLAE, in combination with the pharmacological or genetic manipulations on the above-mentioned memory tests. Therefore, as for the analyses of the two pharmacological approaches, the main factors were defined as PLAE (levels: water and PLAE) and treatment (levels for the first analysis shown in Fig3.B-D: VEH-VEH, GW9662-VEH, VEH-URB597, and GW9662-URB597; levels for the second analysis shown in Fig3.F-H: VEH and PIO). Regarding the results of the viral vector approach, the main factors were defined as PLAE (levels: water and PLAE) and PPAR-γ (levels: AAV5-GFAP-mCherry and AAV5-GFAP-PPAR-γ-mCherry). When significant interactions were observed in the ANOVAs, individual comparisons were performed with Sidak’s post-hoc tests. The statistical significance was set at p<0.05. Outliers were calculated for each analysis using Grubb’s test. Statistical analyses were made with GraphPad Prism 8.0. software.

## Results

### 1. Maternal alcohol drinking during gestation and lactation

Dams were allowed to access either water or alcohol (20%) during both gestation and lactation periods following the DID test schedule (Fig1.A). Liquid volumes were recorded before and after every drinking session. The water and alcohol consumption (volume in ml) over the 6 weeks is presented in Fig1.B. The fluid intake was then calculated for dams exposed to alcohol and is represented in Fig1.C. This is an indirect measure of alcohol consumption by the dams, however, our team has previously demonstrated a correlation between alcohol intakes and moderate-to-high blood alcohol concentration in dams (3) as well as in pups (58).

**Figure 1.**
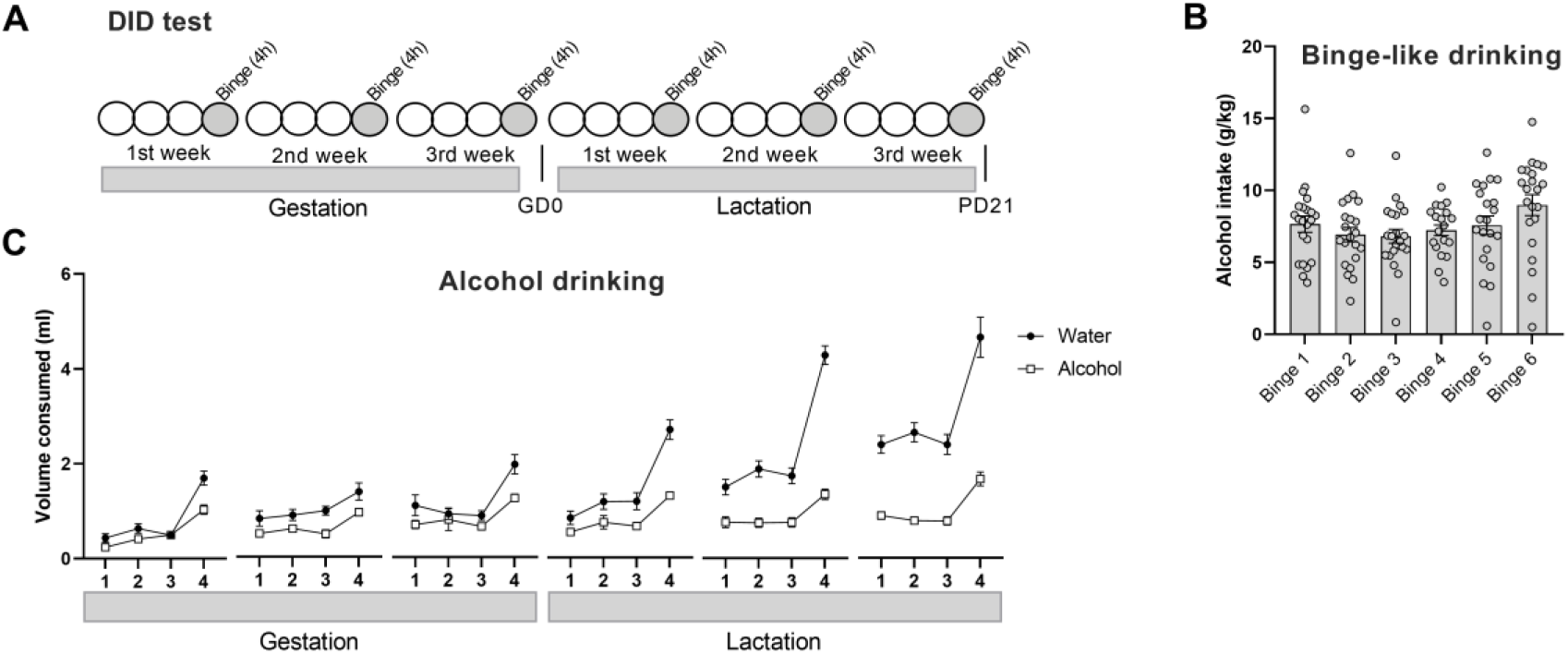
Alcohol consumption by dams during gestation and lactation. A) Schematic representation of the DID test. B) Volumes (ml) of water or alcohol solution consumed by dams during the DID test (N= 24 dams for water; N= 23 dams for alcohol). C) Alcohol intake (g of alcohol per kg of body weight) in the six sessions of binge-like drinking. Data are presented as mean ± SEM. DID, drinking in the dark.

### 2. PLAE reduces the hippocampal levels of NAEs, PPAR-γ and CB1 at an early life stage

We analyzed the levels of NAEs and the expression of genes that belong to the expanded ECS at two developmental times (PD25 and PD70) in both HPC and PFC. At PD25, we found a reduction in the levels of N-oleoylethanolamine (OEA) (Fig2.A; t=2.684, p<0.05), N-palmitoylethanolamine (PEA) (Fig2.A; t=5.094, p<0.001), N-docosatetraenoylethanolamine (DEA) (Fig2.A; t=3.017, p<0.01), N-docosahexaenoylethanolamine (DHEA) (Fig2.A; t=2.34, p<0.05), as well as PPAR-γ (Fig2.B; t=5.055, p<0.001) and CB1 (Fig2.B; t=4.052, p<0.01) expression in the HPC of PLAE when compared to control mice. However, in the PFC, we only found an increase in 2-arachidonoylglycerol (2-AG) levels at this early life stage (FigS1.A-B). However, no differences were found at PD70 in either the HPC (Fig2.C-D) or the PFC (FigS1.C-D).

**Figure 2.**
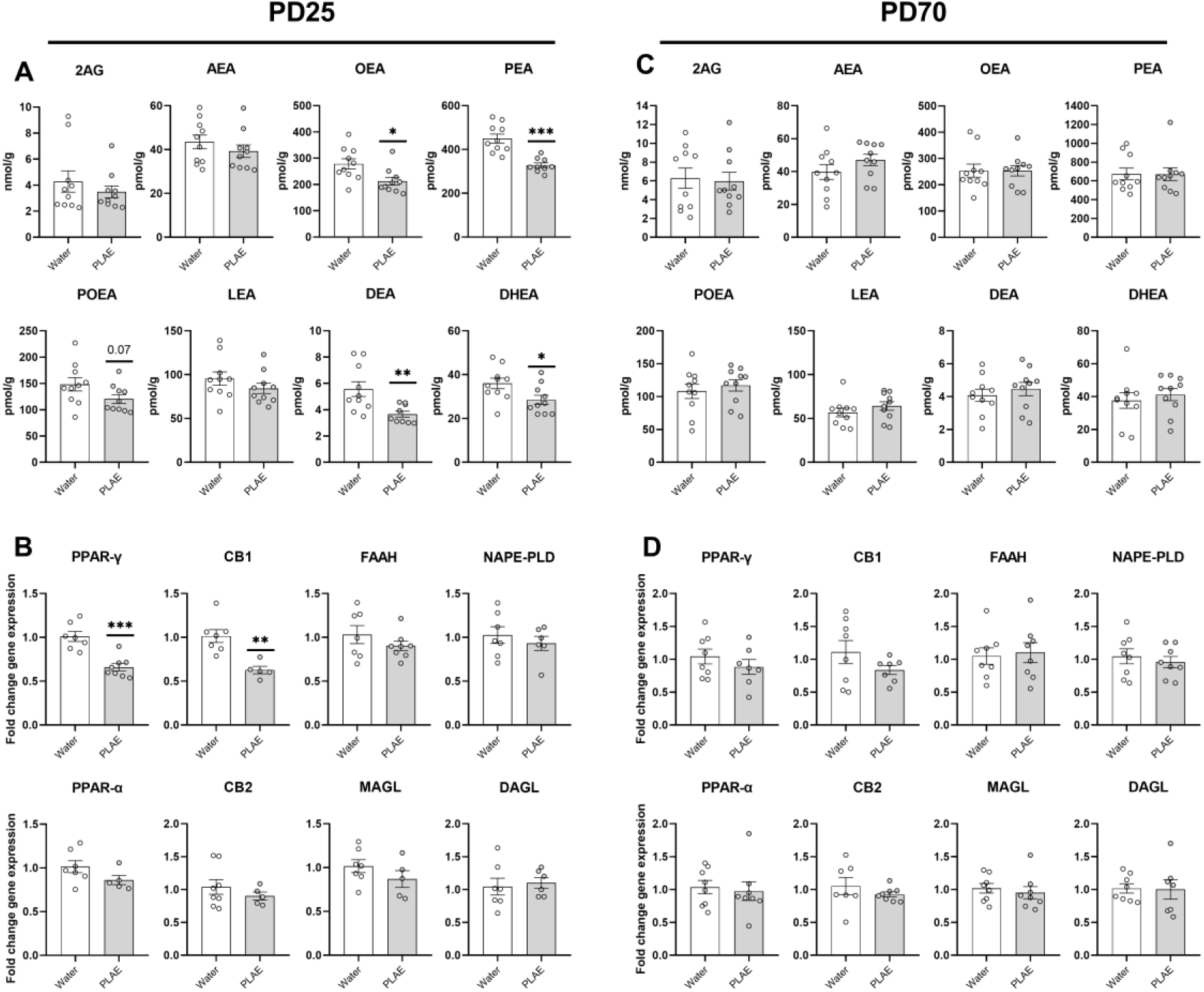
Hippocampal alterations in the expanded ECS induced by PLAE. A) Levels of endocannabinoids and other NAEs (nmol or pmol per grams of tissue) at PD25 (N= 9-10 mice/group) and C) at PD70 (N= 9-10 mice/group) in the HPC. B) Fold change (calculated based on 2^-ΔΔCT) of gene expression of receptors and enzymes related to the expanded ECS at PD25 (N= 5-8 mice/group) and D) at PD70 (N= 7-8 mice/group) in the HPC. Data are presented as mean ± SEM. Significant differences between water and PLAE groups revealed by unpaired t-Tests are represented * p<0.05, ** p<0.01, *** p<0.001. 2-AG, 2-arachidonoylglycerol; AEA, anandamide; DEA, N-docosatetraenoylethanolamine; DHEA, N-docosahexaenoylethanolamine; LEA, N-linoleoylethanolamine; OEA, N-oleoylethanolamine; PEA, N-palmitoylethanolamine; POEA, N-palmitoleoylethanolamine; PPAR-γ, peroxisome proliferator activated receptor type gamma; PPAR-a, peroxisome proliferator activated receptor type alpha; CB1, cannabinoid receptor type 1; CB2, cannabinoid receptor type 2; FAAH fatty acid amide hydrolase; MAGL, monoacylglycerol lipase; NAPE-PLD, N-acyl phosphatidylethanolamine-specific phospholipase D; DAGL, diacylglycerol lipase; PD, post-partum day; PLAE, prenatal and lactation alcohol exposure.

These results suggest that a disruption of the expanded ECS at an early age could be involved in memory deficits observed in adult PLAE mice.

### 3. The pharmacological activation of PPAR-γ at an early life stage rescues memory deficits in PLAE mice

We performed two pharmacological approaches during an early life stage (from PD25 to PD34) to explore whether PPAR-γ activation, either by endogenous NAEs or the PPAR-γ agonist pioglitazone, is involved in memory deficits present in adult PLAE mice (Fig3.A). For that, we administered the FAAH inhibitor URB597 (0.3 mg/kg, i.p.) and the PPAR-γ antagonist GW9662 (1 mg/kg, i.p.) to a set of mice. In another experiment, we administered pioglitazone (10 mg/kg, i.p.) to PLAE and control mice.

**Figure 3.**
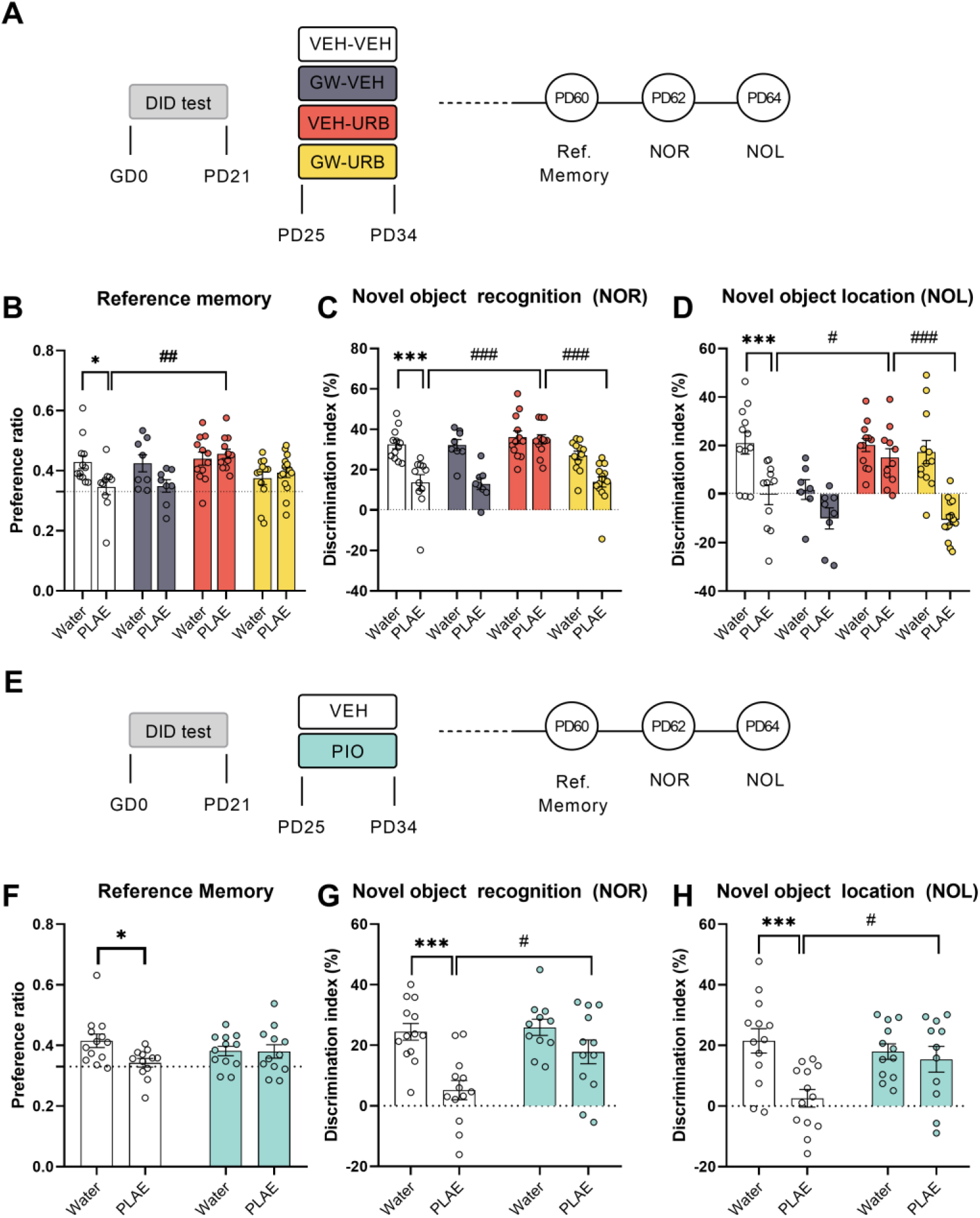
Endogenous or exogenous PPAR-γ activation ameliorates memory impairments in PLAE mice. A, E) Schematic timelines of the two pharmacological approaches: after the weaning, water-exposed and PLAE mice were daily treated from PD25 to PD34. Then, memory tests were evaluated when they became adults at PD60, PD62 and PD64. B) Preference ratio calculated in the reference memory test to evaluate spatial reference memory. The dashed grid line indicates a random preference that would be found if an animal spent equal time in each arm of the Y-maze. Two-way ANOVA, * p<0.05 Water-VEH vs PLAE-VEH, ## p<0.01 PLAE-VEH vs PLAE-URB. C) Percentage of discrimination index calculated in the NOR test to assess object recognition memory. Two-way ANOVA, *** p<0.001 Water-VEH vs PLAE-VEH, ### p<0.001 PLAE-VEH vs PLAE-URB and PLAE-URB vs PLAE-GW-URB. D) Percentage of discrimination index calculated in the NOL test to evaluate object location memory. Two-way ANOVA, *** p<0.001 Water-VEH vs PLAE-VEH, # p<0.05 PLAE-VEH vs PLAE-URB, ### p<0.001 PLAE-URB vs PLAE-GW-URB. (N= water-VEH: 12; PLAE-VEH: 11; water-GW: 8; PLAE-GW: 8; water-URB: 12; PLAE-URB: 12; water-GW-URB: 12; PLAE-GW-URB: 15 mice). F) Preference ratio as in Fig3.B. Two-way ANOVA, * p<0.05 Water-VEH vs PLAE-VEH. G) Percentage of discrimination index in the NOR test as in Fig3.C. Two-way ANOVA, *** p<0.001 Water-VEH vs PLAE-VEH, # p<0.05 PLAE-VEH vs PLAE-PIO. H) Percentage of discrimination index in the NOL test as in Fig3.D. Two-way ANOVA, *** p<0.001 Water-VEH vs PLAE-VEH, # p<0.05 PLAE-VEH vs PLAE-PIO. (N= 11-13 mice/group). DID, drinking in the dark; PLAE, prenatal and lactation alcohol exposure; VEH, vehicle; GW, GW9662; URB, URB597; PIO, pioglitazone; PD, post-partum day; NOR, novel object recognition; NOL, novel object location.

*Post-hoc* analyses revealed that PLAE-VEH-VEH mice show impaired spatial reference (Fig3.B, p<0.05; PLAE effect: F(1,81)= 4.162, p<0.05), object recognition (Fig3.C, p<0.001; PLAE effect: F(1,82)= 43.11, p<0.001) and object location memories (Fig3.D, p<0.001; PLAE effect: F(1,80)= 36.25, p<0.001) compared to water-VEH-VEH mice. As for the object recognition memory, the PLAE-induced impairment was still exhibited when comparing PLAE-GW9662-VEH to water-GW9662-VEH (Fig3.C, p<0.001), and PLAE-GW9662-URB597 to water-GW9662-URB597 (Fig3.C, p<0.001). In case of object location memory, the PLAE effect was still found when PLAE-GW9662-URB597 was compared to water-GW9662-URB597 (Fig3.D, p<0.001).

URB597 treatment improves memory deficits in PLAE mice as *post-hoc* analyses revealed when comparing PLAE-VEH-URB597 to PLAE-VEH-VEH groups in spatial reference (Fig3.B, p<0.01; interaction PLAE x Treatment: F(3,81)= 3.476, p<0.05), object recognition (Fig3.C, p<0.001; interaction: F(3,82)= 4.655, p<0.01), and object location (Fig3.D, p<0.05; interaction: F(3,80)= 3.812, p<0.05) memories. Interestingly, GW9662 prevents the beneficial effects of URB597 on memory tasks when co-administered. *Post-hoc* comparisons revealed statistically significant differences between PLAE-VEH-URB597 and PLAE-GW9662-URB597 in the object recognition (Fig3.C, p<0.001) and the object location (Fig3.D, p<0.001) memories.

In a separated batch of animals, we replicated the memory impairments induced by PLAE as indicated by significant differences found between PLAE-VEH and water-VEH in spatial reference (Fig3.E, p<0.05; PLAE effect: F(1,45)= 3.769, p=0.05), in object recognition (Fig3.F, p<0.001; PLAE effect: F(1,45)= 18.4, p<0.001), and in object location (Fig3.E, p<0.001; PLAE effect: F(1,45)= 9.453, p<0.01) memories. In this batch, pioglitazone (PIO) treatment also attenuates memory deficits in PLAE mice as *post-hoc* analyses indicate after comparing PLAE-VEH to PLAE-PIO in object recognition (Fig3.F, p<0.05; treatment effect: F(1,45)= 4.903, p<0.05) and object location (Fig.G, p<0.05; interaction: F(1,45)= 5.507, p<0.05).

Altogether, these results show that URB597 treatment during development restores PLAE-induced memory deficits through a PPAR-γ dependent mechanism. Moreover, pioglitazone treatment during development also improves memory test performance suggesting a key role of PPAR-γ signaling in memory deficits in PLAE mice.

### 4. PLAE reduces PPAR-γ in hippocampal astrocytes

To further explore which cell types show PLAE-induced reductions of PPAR-γ in the HPC, we performed immunohistochemical analyses to co-localize PPAR-γ with GFAP or NeuN at PD25. Interestingly, our data exhibited that PPAR-γ labelling can show two different shapes depending on neuronal or astrocytic location (FigS2.A), as it has been previously showed (47).

Unpaired t-test revealed that PLAE mice show fewer PPAR-γ positive cells (Fig4.A; t= 2.629, p<0.05) and PPAR-γ/GFAP positive cells (Fig4.B; t= 2.499, p<0.05) in hippocampal CA1. By contrast, no differences were found in PPAR-γ/NeuN positive cells between groups (FigS2.C-E). Therefore, these findings indicate that PPAR-γ in astrocytes is more sensitive to the effects of early alcohol exposure than it is in neurons and, in turn, astrocytic PPAR-γ might be involved in memory dysfunction induced by PLAE.

**Figure 4.**
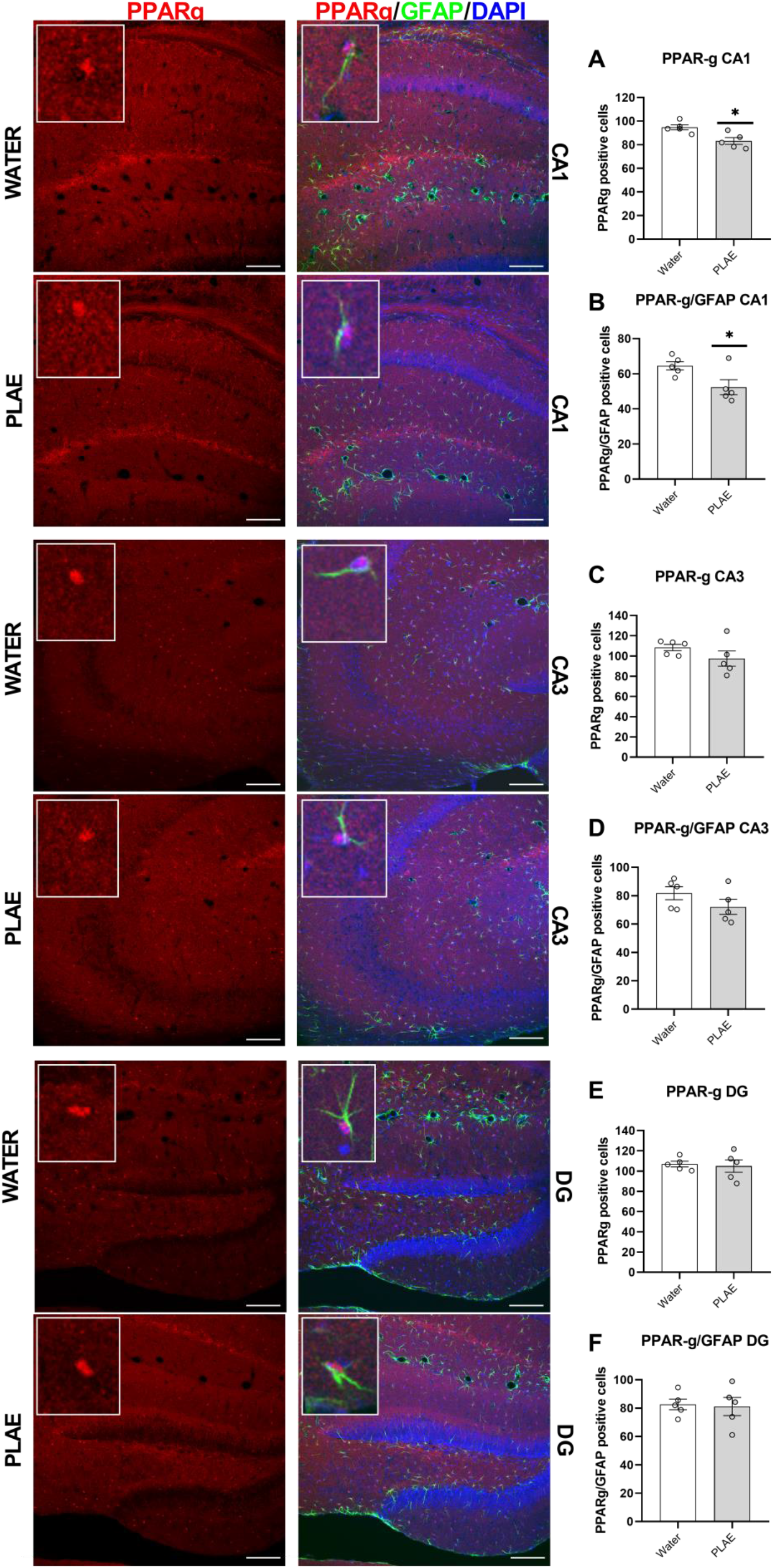
PLAE reduces astrocytic PPAR-γ primarily in CA1 of the HPC. Representative pictures of immunohistochemical analyses of the three main areas of the HPC (CA1, CA3 and DG), showing PPAR-γ positive cells in red, GFAP positive cells in green and DAPI in blue. Total number of PPAR-γ positive cells in A) CA1, C) CA3, E) DG. Total number of PPAR-γ and GFAP (co-localization of the two markers) positive cells in B) CA1, D) CA3, F) DG. (N= 5 mice/group). Data are presented as mean ± SEM. Significant differences between water and PLAE groups revealed by unpaired t-Tests are represented * p<0.05. PPAR-γ, peroxisome proliferator activated receptor type gamma; GFAP, glial fibrillary acidic protein; DG, dentate gyrus: PLAE, prenatal and lactation alcohol exposure. Scale bar 100 μm.

### 5. Overexpression of PPAR-γ in hippocampal astrocytes ameliorates memory deficits in PLAE mice

To unravel whether increasing the expression of PPAR-γ in hippocampal astrocytes is sufficient to restore memory deficits in PLAE mice, we infused a viral vector expressing PPAR-γ under the GFAP promoter around PD25 (Fig5.C). We have validated our viral vector approach by two methods performed in the same mice that underwent the behavioral experiments. On the one hand, we performed histological analyses that showed mCherry staining at the proper coordinates in the two hemispheres (AP=from -1.7 to -2.8 with the highest intensity at -2.18. Fig5.B). On the other hand, in another group of mice, we carried out a RT-qPCR that showed a higher expression of PPAR-γ in animals that received the viral vector containing PPAR-γ (Fig5.D; t=6.878, p<0.001). In total, three mice were removed from the experiment for showing very low intensity of mCherry fluorescence.

**Figure 5.**
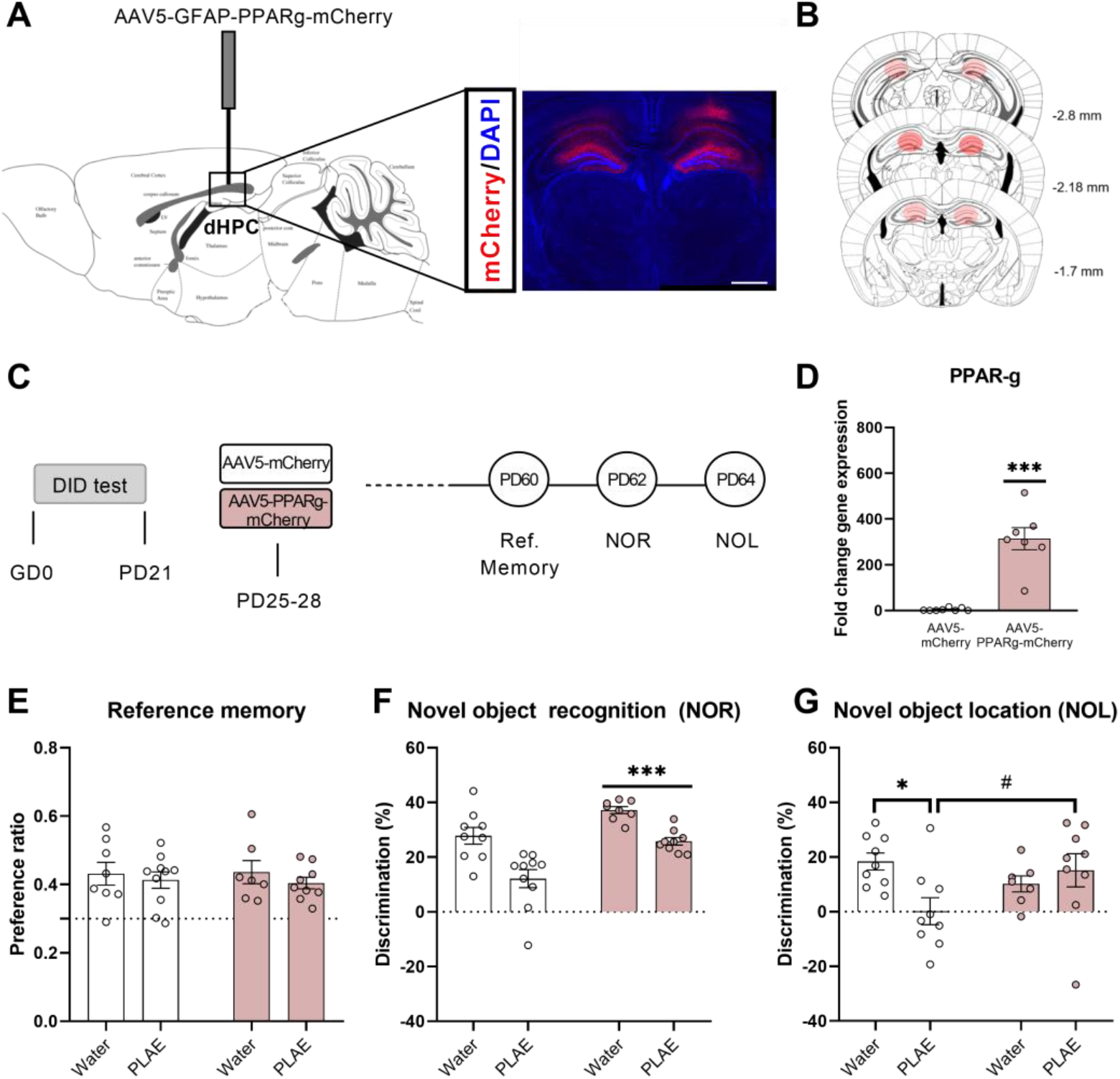
Astrocytic PPAR-γ upregulation in dorsal HPC rescues memory deficits induced by PLAE. A) Strategy used to overexpress PPAR-γ in astrocytes of dorsal HPC and representative immunofluorescence showing mCherry 5 weeks after the injection. AAV5-GFAP-PPARγ-mCherry was infused around PD25. B) Schematic illustration of the injection site and the spread of the viral vector. Note that the spread (determined by mCherry immunofluorescence) was within the AP coordinates -1.7 mm and -2.8 mm. C) Schematic timeline of the experiment: after the weaning, the viral vector (PPARγ-containing or the control) was infused in water-exposed and PLAE mice around PD25. Then, memory tests were evaluated when they became adults at PD60, PD62 and PD64. D) Fold change (calculated based on 2^-ΔΔCT) of PPAR-γ expression in animals that received control or PPARγ-containing viral vector. E) Preference ratio calculated in the reference memory test to evaluate spatial reference memory. The dashed grid line indicates a random preference that would be found if an animal spent equal time in each arm of the Y-maze. F) Percentage of discrimination index calculated in the NOR test to assess object recognition memory. Two-way ANOVA, *** p<0.001 PPAR-γ effect, meaning that animals that received the PPARγ-containing viral vector showed higher discrimination index regardless PLAE factor. G) Percentage of discrimination index calculated in the NOL test to evaluate object location memory. Two-way ANOVA, * p<0.05 WATER-AAV5-mCherry vs PLAE-AAV5-mCherry, # p<0.05 PLAE-AAV5-mCherry vs PLAE-AAV5-PPARγ-mCherry. (N= 7-10 mice/group). dHPC, dorsal hippocampus; DID, drinking in the dark; AAV5-mCherry, AAV5-GFAP-mCherry; AAV5-PPARγ-mCherry, AAV5-GFAP-PPARγ-mCherry; PPAR-γ, peroxisome proliferator activated receptor type gamma; GFAP, glial fibrillary acidic protein; PD, post-partum day; NOR, novel object recognition; NOL, novel object location; PLAE, prenatal and lactation alcohol exposure.

Regarding behavioral data, a two-way ANOVA revealed that PLAE impairs object recognition memory, while PPAR-γ overexpression improves it regardless of the PLAE factor (Fig5.F; PLAE effect: F(1,32)= 27.93, p<0.001; PPAR-γ effect: F(1,32)= 20.27). As for the object location memory, *post-hoc* analyses showed that PLAE mice infused with the control AAV5 still exhibit the impairment compared to its respective water-exposed control group (Fig5.G, p<0.05; interaction: F(1,30)= 6.156, p<0.05). Furthermore, the infusion of AAV5 expressing PPAR-γ attenuates the deficit in object location memory as shown when compared to PLAE mice infused with the control AAV5 (Fig5.G, p<0.05). Lastly, no significant differences were found in reference memory test under these conditions (Fig5.E).

Altogether, this data proves that upregulating astrocytic PPAR-γ in HPC is sufficient to attenuate PLAE-induced memory impairments.

## Discussion

In this study we have identified PPAR-γ activation as a promising strategy to counteract the memory impairments induced by alcohol exposure during pregnancy and lactation. With a combination of behavioral, pharmacological, and genetic approaches we thereby demonstrate that endogenous lipid mediators, a synthetic agonist, or a genetic upregulation of PPAR-γ rescues memory deficits in a mouse model of FASD.

First, using a validated PLAE mouse model (3,9), we report that persistent memory impairments are accompanied by alterations in the expanded ECS. At an early life stage (PD25), we observed a reduction of CB1 and PPAR-γ expression, as well as reduced levels of OEA, PEA, DEA and DHEA in the HPC of PLAE mice. In contrast to this strong hippocampal impact, we only found an increase in 2-AG in the PFC. In line with these findings, other authors have reported that perinatal alcohol exposure affects brain development via an ECS-related mechanism that contributes to the long-lasting learning and memory deficits in rodents (59). However, they described increased levels of AEA and CB1 in the neocortex and the HPC throughout the 24 h following an acute alcohol administration at PD7, while no changes were observed in 2-AG (25). The inconsistencies between both studies might be explained by the protocol used for alcohol exposure and/or the timing of the analyses. In our approach, mice were intermittently exposed to alcohol throughout gestation and lactation, whereas in this study (25) there was only an acute exposure at PD7. Moreover, we observed that the effects of early alcohol exposure on the expanded ECS are time-dependent. In fact, we report significant alterations earlier in life (around PD25) that disappear during adulthood, since no differences were found at PD70. Interestingly, our results suggest a link between reduced levels of NAEs during neurodevelopment and persistent memory deficits in PLAE mice. We show that increasing the levels of NAEs via peripheral administration of the FAAH inhibitor URB597 from PD25 to PD34 restores memory deficits, which supports that early life stages represent a promising therapeutic window. NAEs apart from AEA have low to no affinity for the canonical ECS receptors CB1 and CB2 thus, their biological function is often mediated by other receptors like nuclear PPARs (15,60). Importantly, the neuroprotective role of said NAEs-PPARs pathway, as well as PPARs activation by exogenous cannabinoids, in the context of cognitive dysfunction has been widely described (15,22). In line with this literature, we demonstrate that the beneficial effects of URB597 treatment on PLAE-induced memory impairments are mediated by PPAR-γ. In our approach, the co-administration of the selective PPAR-γ antagonist GW9662 suppresses the beneficial effects of URB597 on memory dysfunction. Therefore, our data provide proof of the neuroprotective role of PPAR-γ activation by endogenous NAEs in PLAE mice.

In a parallel line, it has been observed that early life adverse events alter PPARs expression and function which negatively impacts on health and behavior later in life (18). Particularly, decreased expression of PPAR-γ in the HPC has been correlated with cognitive impairments in preclinical models of neurodevelopmental disorders. In these experimental models, the PPAR-γ agonist pioglitazone improved cognitive impairments, likely due to its well described antioxidant, anti-inflammatory and pro-neurogenic effects (20,21). Moreover, cognitive impairments induced by perinatal alcohol exposure were rescued by natural PPAR-γ ligands, as resveratrol (29) and cannabidiol (9,26). Accordingly, our results show that PLAE downregulates PPAR-γ in the HPC. In addition, the administration of pioglitazone attenuates persistent memory impairments induced by early alcohol exposure. Thus, the two pharmacological approaches point out that either endogenous or exogenous PPAR-γ activation (via increased levels of NAEs or pioglitazone, respectively) during a crucial developmental period rescues memory deficits in PLAE mice.

Lastly, since both the main alterations in the expanded ECS and the behavioral traits rescued by PPAR-γ activation are classically related to hippocampal function (61,62), we studied in depth the cell type and sub-region specificities in PLAE-induced effects on PPAR-γ. Hippocampal PPAR-γ expression is known to be age and brain region-dependent (46). As for cell-type specificity, PPAR-γ has been mainly identified in neurons and astrocytes of adult mice and humans (46,47). In this line, our results show that, at PD25, hippocampal PPAR-γ is found in both neurons and astrocytes. Nonetheless, in accordance with the PPAR-γ downregulation reported by q-PCR, semi-quantitative immunohistochemistry analyses indicated that PLAE reduces PPAR-γ levels specifically in astrocytes primarily located in hippocampal CA1.

Accordingly, prominent alterations in astrocytes development and function have been described following alcohol exposure *in vitro* and *in vivo* (63–65), and appropriate astrocytic function in the HPC is key for memory encoding and recall (38). To evaluate the role of astrocytic PPAR-γ in PLAE, we increased PPAR-γ expression specifically in astrocytes in the dorsal HPC using a viral approach. Our manipulation was sufficient to rescue persistent memory deficits in PLAE mice showing a link between reduced expression of astrocytic PPAR-γ during neurodevelopment and cognitive dysfunction induced by early alcohol exposure. Unfortunately, due to the variety and abundance of PPAR-γ targets, the exact downstream mechanisms by which its overexpression restores PLAE deficits will remain, for the moment, unclear. However, a recent study in which authors addressed a similar neurodevelopmental period reported that the downregulation of the PPAR-γ coactivator 1 alpha (PGC-1alpha) disrupts mitochondrial biogenesis in developing astrocytes, impairing astrocytic maturation and synaptogenesis (66). In fact, extensive evidence indicates that PPAR-γ is involved in mitochondrial biogenesis partly due to its interaction with PGC-1alpha (67,68). Hence, one could hypothesize that restoring astrocytic mitochondrial biogenesis might represent a putative mechanism explaining how overexpressing PPAR-γ in hippocampal astrocytes during development restores memory deficits in PLAE mice. Despite that, alternative mechanisms in which astrocytic PPAR-γ is involved, such as regulation of neuroinflammation (69,70) or brain energy homeostasis through glucose metabolism (71), cannot be discarded. Indeed, some authors have suggested that PPAR-γ agonists may prevent neurotoxicity in FASD through an anti-inflammatory mechanism dependent on microglia and astrocytes (10).

In summary, our findings provide strong evidence that PPAR-γ signaling during a key temporal window of neurodevelopment is involved in PLAE-induced memory deficits. Indeed, systemic PPAR-γ activation, by both an exogenous agonist and endogenous NAEs, ameliorated spatial, recognition and location memory impairments. As to the exact mechanisms by which PPAR-γ activity is exerting these beneficial effects on PLAE, our findings indicate that targeting PPAR-γ expression in hippocampal astrocytes is an effective tool to rescue memory dysfunction. Therefore, these findings represent an essential key in the understanding of cognitive disfunction related to memory in FASD patients. This is of outmost importance since cognitive deficits in FASD are highly correlated with the patients’ functional disabilities (59,60) and, therefore, effective pharmacological treatments could significantly improve these patients’ quality of life. Hence, we conclude that further understanding of astrocytic PPAR-γ function might be useful to design promising future treatments for memory impairments in FASD.

## Acknowledgments

This work was supported by the Ministerio de Economia y Competitividad (grant number PID2019-104077RB-100), Ministerio de Sanidad (Retic-ISCIII, RD16/017/010 and Plan Nacional sobre Drogas 2018/007). A.G-B received a FI-AGAUR grant from the Generalitat de Catalunya (grant number 2019FI_B0081). I.G-L. obtained a grant from the Ministerio de Ciencia e Innovación (PRE2020-091923) The Department of Medicine and Health Sciences (UPF) is a “Unidad de Excelencia María de Maeztu” funded by the AEI (grant number CEX2018-000792-M).

The authors wish to thank Xavier Puig-Reyne for the technical support.

