## Supplemental Materials for "PPAR-γ is a promising therapeutic target for memory deficits induced by early alcohol exposure"

*Garcia-Baos A et al.*

### Supplemental material and methods

#### 1. Drinking in the Dark (DID) test

Two days after mating, pregnant females were randomly assigned to two groups: alcohol or water (control). Three hours after the lights were turned off, the water bottles were replaced with 10-ml graduated cylinders fitted with sipper tubes containing either 20% (v/v) alcohol in tap water or only tap water. From Monday to Wednesday (days 1, 2, 3), pregnant females were allowed to voluntarily drink for 2 h-access period. On Thursday (day 4), the drinking-access period was extended to 4 h. Volumes consumed were recorded just after the drinking-access periods. During this procedure, all the females were individually housed to record individual fluid intakes. Following the drinking-access period, regular water bottles were returned to the home cage. Fluid intakes (g/kg of body weight) were calculated for each day on the basis of average 2-day body weight values, since dams were weighed at 2-day intervals (Monday and Wednesday).

In our experiment, DID test consisted of six consecutive weeks, encompassing prenatal and lactation periods. Therefore, each pregnant dam was exposed to six binge-like drinking sessions (day 4), following the habituation period (days 1,2,3). As previously reported, blood alcohol concentration of dams reached levels of ~0.8 g/L after the last binge-like drinking session (1), which is in accordance with “Drinking Levels Defined” reported by NIAAA (2).

### 2. Quantitative reverse transcription polymerase chain reaction (RT-qPCR)

Water-exposed and prenatal and lactation alcohol exposed (PLAE) animals were euthanized by cervical dislocation at post-partum day (PD) 25 and PD70, both in basal conditions. Fresh hippocampus (HPC) and prefrontal cortex (PFC) tissue were dissected and stored at -80ºC until the molecular analyses. As for the RT-qPCR, we first extract the RNA from the samples using Trizol (ThermoFisher Scientific) as previously described (3). Retrotranscription was performed using High-Capacity cDNA Reverse Transcription Kit (ThermoFisher Scientific) following the manufacturer's protocol. SYBR Green Master Mix (ThermoFisher Scientific), 100 ng of cDNA and the respective primers at 0.2 µM (Table S1) were used for the RT-qPCR. The qPCR was performed in QuantStudio 12K Flex (ThermoFisher) using the following program: 95 °C-10s, 60 °C-20s, 72 °C-10s for 40 cycles. The threshold cycle (CT) for each target product was determined and the ΔΔCT method was used to calculate the fold change relative to the control group in each case. The CT of each target product was normalized to housekeeping gene GAPDH. The CT values of the housekeeping gene were not significantly different among the experimental groups.

### 3. Quantification of endocannabinoids and other related compounds by liquid chromatography–tandem mass spectrometry (LC-MS/MS)

We measured the levels of endocannabinoids and other related compounds in two brain areas (PFC and HPC) firmly vulnerable to alcohol effects and involved in memory performance. Animals underwent the DID test were euthanized and samples were collected at PD25 and PD70 as described in the previous section.

The quantification of these compounds was done as previously described (4). The following compounds were measured: 2-arachidonoylglycerol (2-AG), N-arachidonoylethanolamine or anandamide (AEA), N-docosatetraenoylethanolamine (DEA), N-docosahexaenoylethanolamine (DHEA), N-linoleoylethanolamine (LEA), N-oleoylethanolamine (OEA), N-palmitoylethanolamine (PEA), N-palmitoleoylethanolamine (POEA). Brain tissue was homogenized on ice with a glass homogenizer using a mixture of Tris-HCl buffer 50 mM (pH 7.4) and methanol (1:1) and extracted with chloroform. The extracts were analysed by LC-MS/MS with an Agilent 6410 triple quadrupole (Agilent Technologies, Wilmington, DE), equipped with a 1200 series binary pump, a column oven and a cooled autosampler (4 °C). Chromatographic separation was carried out by gradient chromatography using an ACQUITY UPLC BEH C18 column (3.1 × 100 mm, 1.8 μm particle size) maintained at 40 °C with a mobile phase of water/acetonitrile containing 0.1% formic acid, and a flow rate of 0.4 ml/min. Quantification was done by isotope dilution with the response of the internal standards from Cayman Chemical (Ann Harbor, MI, USA): 2-AG-d5, AEA-d4, DHEA-d4, LEA-d4, OEA-d4, and PEA-d4. The quantification of 2-AG was done as the sum of the two isomers 1-AG and 2-AG.

### 4. Immunohistochemistry

Water-exposed and PLAE mice at PD25 were deeply anesthetized using a Dolethal overdose (i.p. injection of 120 mg/kg of pentobarbital-based solution). Then, they underwent a transcardiac perfusion of phosphate saline solution 0.1M (PBS, pH 7.4), followed by 4% paraformaldehyde in 0.1M phosphate buffer pH 7.4 using a peristaltic pump (5.5ml/min for 2 and 5 min, respectively). Brains were post-fixed in the same fixative solution overnight at 4 °C. Then, brains were placed into 30% sucrose solution (in 0.1 M PBS, pH 7.6, 4 °C) until they sank. Finally, the brains were frozen, and we obtained 40-μm-thick coronal sections with a cryostat (Leica CM3050 S, Wetzlar, Germany). Free-floating sections were collected in five parallel sets.

Two out of five parallel sets were used for the experiment: i) colabelling of PPAR-γ and NeuN (marker of neuronal nuclei), ii) colabelling of PPAR-γ and GFAP (marker of astrocytes), to explore the effects of PLAE on neuronal and astrocytic PPAR-γ in the HPC. Briefly, sections underwent an antigen retrieval step to allow PPAR-γ labelling: first, they were incubated for 20 min in 37% chloridric acid (HCl) 2N in distilled water at 37 ºC, followed by an incubation in a basic solution of borate buffer 0.1M pH 8.5 for 10 min to quenched the HCl. Afterwards, sections were pre-incubated in blocking solution of 4% normal goat serum in 0.05M TRIS buffer saline (TBS) pH 7.4 with 0.2% Triton X-100 at RT for 1h. Then, sections were incubated in primary antibodies (overnight at 4 ºC) diluted in blocking solution: rabbit anti-PPAR-γ (1:100, Cell signalling, #C26H12), mouse anti-GFAP conjugated with Alexa Fluor 488 (1:1000, ThermoFisher Scientific, #53-9892-82), or mouse anti-NeuN (1:1000, Merck Millipore, #MAB377). Then, they were incubated with fluorescent-conjugated secondary antibodies (90 min at RT) diluted in blocking solution: Alexa Fluor 488-conjugated goat anti-rabbit IgG (1:1000, ThermoFisher Scientific, #A11070), Alexa Fluor 555-conjugated goat anti-mouse IgG (1:1000, ThermoFisher Scientific, #A-21422), Alexa Fluor 555-conjugated goat anti-rabbit IgG (1:1000, ThermoFisher Scientific, #A-21428). The tissue was washed in TBS in between each step (except after blocking solution). Before mounting, sections were washed with TB to prevent crystal formation. Finally, they were mounted onto gelatinized slides and cover-slipped with fluorescence mounting medium containing DAPI (Fluoromount-G$^{TM}$, ThermoFisher Scientific, # 00-4959-52) to reveal the cytoarchitecture of the brain.

**5. Immunofluorescence analysis**

Triple scans were performed to obtain images containing labelling of PPAR-γ, GFAP or NeuN, and DAPI, using a confocal microscope LEICA TCS SP5 upright (Leica, Wetzlar, Germany). Planes with a 2.5 µm Z distance of separation were taken from the region of interest. A stack of 7 Z planes including the 3 channels were taken per image. To minimize the channel spillover, images were sequentially acquired at 20x magnification and saved as LIFF files. For each mouse, we took images of three hippocampal regions, that are CA1, CA3 and DG, of both hemispheres at two different anteroposterior coordinates (Bregma -2 mm and -2.3 mm) to have a greater hippocampal representation. The stacks obtained were processed and analyzed with Fiji-Image J software (#SCR_003070, NIH, Bethesda, MD, USA). Four stacks were analyzed per hippocampal region (two hemispheres of two slices with different anteroposterior coordinates). Three consecutive planes of every stack were analyzed. First, PPAR-γ positive nuclei were manually counted with the counter plugin of Image J software. To prove that PPAR-γ was in nuclei, DAPI colocalization was used when counting. Then, colocalization of PPAR-γ with GFAP or NeuN positive cells was manually counted in the same plane right after to see whether PPAR-γ is expressed by astrocytes or neurons, respectively.

To quantify the total number of PPAR-γ positive cells and PPAR-γ/GFAP or PPAR-γ/NeuN colocalization, we used the following rules for each mouse: i) first, the mean of three consecutive planes per stack was calculated, ii) second, the mean of the two hemispheres per slice was calculated, iii) finally, the mean of the two slices (at two different anteroposterior coordinates) was calculated to plot a greater representation of PPAR-γ expression in each region of the HPC. The analyses were carried out by a blind researcher to the experimental groups.

### 6. Viral vector generation and delivery

The viral vectors AAV5-GFAP-PPARγ-IRES-mCherry-oPRE and its control AAV5-GFAP-IRES-mCherry-oPRE were designed in the free platform of the company VectorBuilder, Inc (Chicago, IL, USA). The plasmid constructs and the viral vectors production were generated by VectorBuilder as well. The inserted PPAR-γ gene was mouse-specific according to the database of PubMed. A 1518 bp fragment of the PPAR-γ gene was inserted into the plasmid vector (residues 46-1563 of mus musculus chromosome 6; accession NM_011146.3).

For viral vector delivery, mice were deeply anesthetized with ketamine hydrochloride (75mg/kg, Imalgene1000, Lyon, France) and medetomidine hydrochloride (1mg/kg, Medeson®, Barcelona, Spain) in a volume of 0.1ml/10g of body weight i.p. Then, after receiving meloxicam (0.5 mg/kg s.c.; Metacam®, Barcelona, Spain) as analgesia, they were placed on the stereotaxic frame (Kopf Instruments, Tujunga, CA). Body temperature was maintained at 37 ºC using a heating blanket. Mice from PD25-PD28 were bilaterally infused in the following coordinates (HPC: -2.2mm anteroposterior; ± 1.7mm mediolateral; -1.7mm dorsoventral from Bregma, 0º angle). 0.3µl/hemisphere of viral vector was manually infused (0.1µl was infused every 5 minutes) using a 5µl Hamilton syringe 75 SN attached to a 33-gauge needle (Teknokroma Analitica S.A., Spain, #HA-87908). The syringe was left in place for an additional 5 minutes to permit diffusion. At the end of the surgery, animals received an injection of atipamezole hydrochloride (0.5mg/kg i.p.; Revertor®, Barcelona, Spain) and 0.1ml glucose

5% solution. Home cages were placed on heating blankets to avoid post-anesthesia hypothermia for 24h. Behavioral experiments began 4 weeks after the surgery when mice were at PD60.

#### 7. Verification and characterization of AAV-expressing PPAR-γ gene

To verify viral vector placements, the same procedure as in the section 5 of supplementary materials and methods was performed with minor modifications. In this case, hippocampal slices were incubated in Hoechst (1:10000; Invitrogen, #33258) for 5 minutes at RT. Then they were washed 2x in TBS 0.05M and once in PB 0.05M before mounting. Images of the HPC at 10x magnification were taken to map the viral vector expression using mCherry fluorescent reporter. Mice either not showing bilateral AAV expression or expression in other areas were excluded from all the analysis.

With another batch of animals (n=4/group) we carried out a functional characterization of the viral vector. Fresh hippocampal tissue was removed after euthanasia by cervical dislocation. Then, mRNA of the samples was isolated and RT-qPCR (as described in the above section 3) was performed to determine whether infusion of AAV5-expressing PPAR-γ overexpresses PPAR-γ in the HPC compared to the respective control.

All these analyses were performed at PD70 (>4 weeks after the viral injections).

#### 8. Behavioral tests for memory

Mice were allowed to acclimatize to the new environmental conditions for at least 1 week prior to experimentation, which occurred during the dark phase under a dim red light. Weeks after the pharmacological or genetic manipulations, the offspring underwent a battery of behavioral tests. Reference memory test was performed at PD60, followed by NOR at PD62 and NOL at PD64.

#### 9.1. Reference memory test

A black Y-maze (three equal 395 mm-long arms, separated by 120º angles) was employed to assess short-term spatial reference memory with 1 h of inter-trial interval, as previously described by our laboratory (5). The time spent in each of the three arms was measured by the Smart Software (Panlab S.L.U., Barcelona, Spain). The preference ratio was calculated by: $\frac{\text{t novel arm}}{\text{t total}}$ , being “t” the time each mouse spent exploring the arms.

#### 9.2. Novel Object Recognition (NOR) test

Black open boxes (24 cm × 24 cm × 15 cm) and plastic toys of similar size to mice were used in this experiment to assess object recognition memory. As previously described by our laboratory (5), this task consists of 3 phases: habituation to the boxes for 5 min, familiarization to two identical objects for 10 min, and the test phase to identify the novel object for 10 min (4 h after the familiarization). To evaluate the memory performance, a discrimination index was calculated as: $\frac{\text{t novel} - \text{t familiar}}{\text{t novel} + \text{t familiar}}\ x\ 100$ , being “t” the time a mouse spent exploring the objects (manually recorded from a video by an observer who was blind to the experimental groups).

#### 9.3. Novel Object Location (NOL) test

This test was performed under the same conditions as NOR as previously described by our laboratory (5), but every mouse was exposed to different objects from the ones used in NOR. In this case, the 3 phases are: habituation for 5 min, familiarization to the location of two identical objects for 10 min, and the test phase to identify the novel location of one of the objects for 10 min. Then, the discrimination index calculated in this case was: $\frac{\text{t displaced object} - \text{t non displaced object}}{\text{t displaced object} + \text{t non displaced object}}\ x\ 100$, being "t" the time a mouse spent exploring the objects (manually recorded from a video by an observer who was blind to the experimental groups).

### Supplemental Results

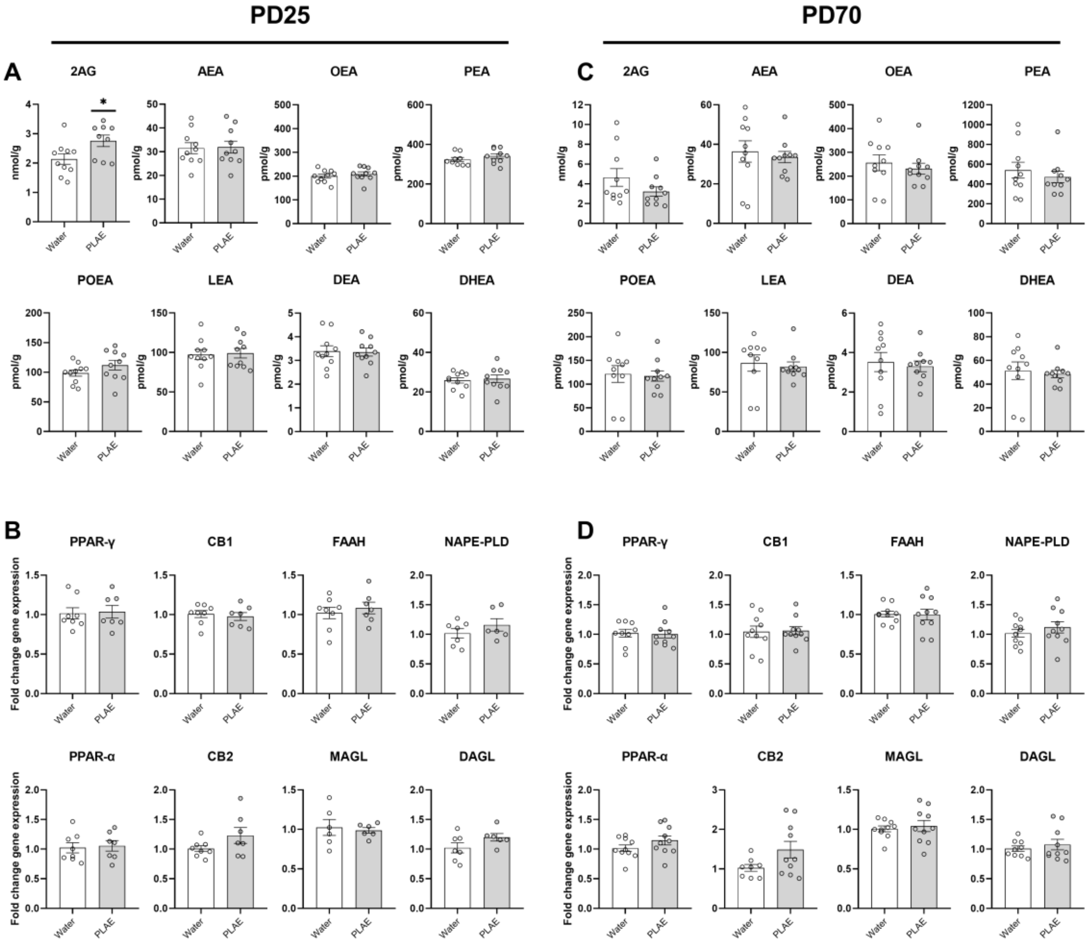


**Figure S1. Alterations in the expanded ECS in the PFC induced by PLAE.**

A) Levels of endocannabinoids and other related lipid compounds at PD25 (N= 9-10 mice/group) and C) at PD70 (N= 9-10 mice/group) in the PFC. B) Fold change (calculated based on 2^-ΔΔCT) of gene expression of receptors and enzymes related to the expanded ECS at PD25 (N= 6-8 mice/group) and D) at PD70 (N= 8-10 mice/group) in the PFC. Data are presented as mean ± SEM. Significant differences between water and PLAE groups revealed by unpaired t-Tests are represented * $p<0.05$. 2-AG, 2-arachidonoylglycerol; AEA, anandamide; DEA, N-docosatetraenoylethanolamine; DHEA, N-docosahexaenoylethanolamine; LEA, N-linoleoylethanolamine; OEA, N-oleoylethanolamine; PEA, N-palmitoylethanolamine; POEA, N-palmitoleoylethanolamine; PPAR-γ, peroxisome proliferator activated receptor type gamma; PPAR-a, peroxisome proliferator activated receptor type alpha; CB1, cannabinoid receptor type 1; CB2, cannabinoid receptor type 2; FAAH fatty acid amide hydrolase; MAGL, monoacylglycerol lipase; NAPE-PLD, N-acyl phosphatidylethanolamine-specific phospholipase D; DAGL, diacylglycerol lipase; PD, post-partum day; PLAE, prenatal and lactation alcohol exposure.

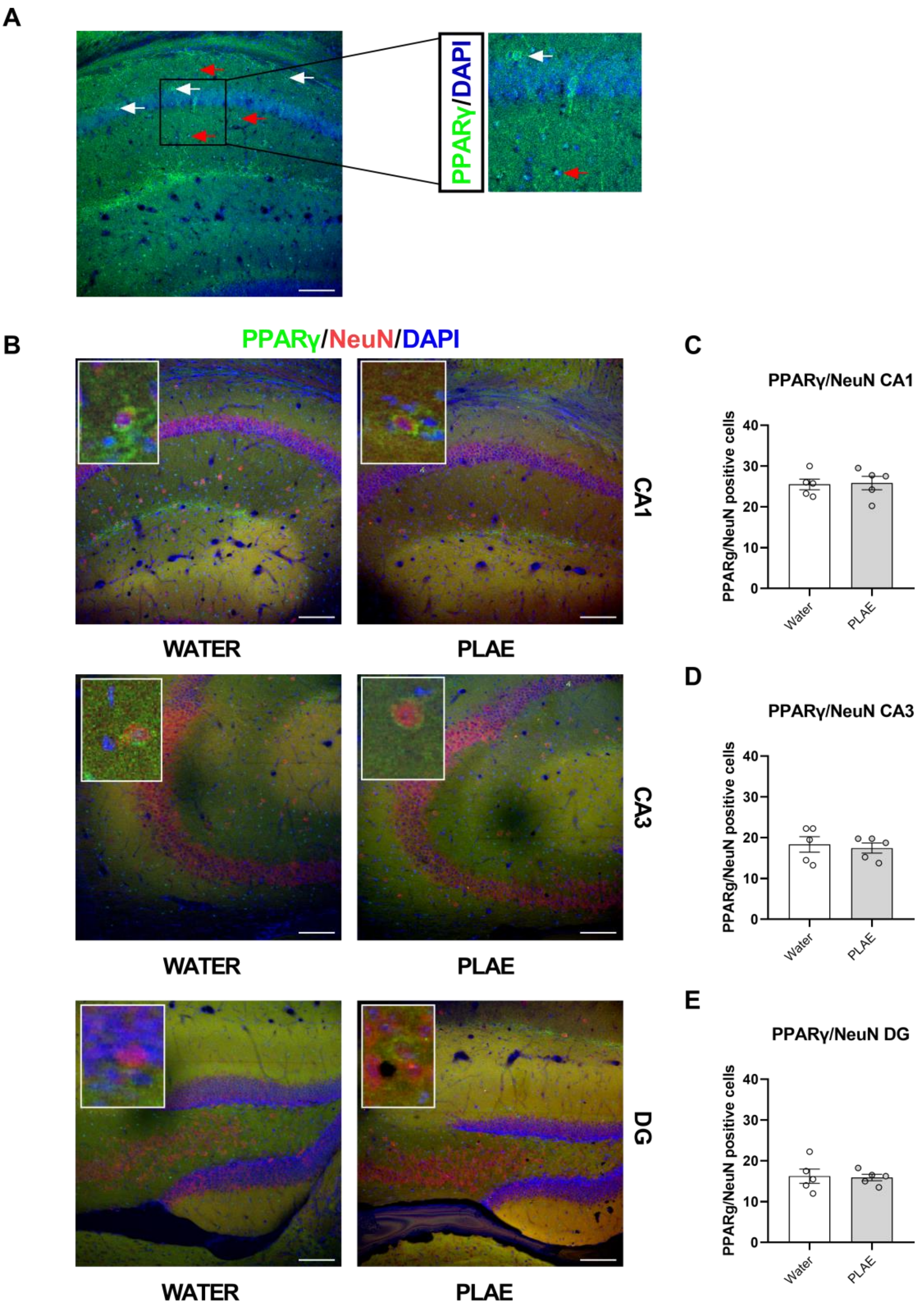


**Figure S2. PLAE does not modify the content of PPAR-γ in hippocampal neurons.** A) Representative picture of PPAR-γ labelling in the CA1, showing two different shapes of PPAR-γ depending on neuronal (white arrows) or astrocytic (red arrows) location. B) Representative pictures of immunohistochemical analyses of the three main areas of the HPC (CA1, CA3 and DG), showing PPAR-γ positive cells in green, NeuN positive cells in red and DAPI in blue. Total number of PPAR-γ and NeuN (co-localization of the two markers) positive cells in C) CA1, D) CA3, E) DG. (N= 5 mice/group). Data are presented as mean ± SEM. PPAR-γ, peroxisome proliferator activated receptor type gamma; NeuN, neuronal nuclear protein; DG, dentate gyrus: PLAE, prenatal and lactation alcohol exposure. Scale bar 100 μm.

Table S1. Primers to quantify gene expression

| | Primer sequence (5'-3') |
|---|---|
| PPAR gamma Forward | CAG GCT TCC ACT ATG GAG TTC |
| PPAR gamma Reverse | GGC AGT TAA GAT CAC ACC TAT CA |
| PPAR alpha Forward | CTG TCG GGA TGT CAC ACA AT |
| PPAR alpha Reverse | CAG GTC GTG TTC ACA GGT AAG |
| CB1R Forward | AGG AGA CAC AAC CAA CAT TAC A |
| CB1R Reverse | TGA AGC ACT CCA TGT CCA TAA A |
| CB2R Forward | GGG TCC TCT CAG CAT TGA TTT |
| CB2R Reverse | AGC CCA GTA GGT AGT CGT TAG |
| NAPE-PLD Forward | CAG ACT AGA GGA GGA CGT AAC T |
| NAPE-PLD Reverse | TCA GCC ATC TGA GCA CAT TC |
| FAAH Forward | GGC TAT CAG CTA CAC TGT TCT C |
| FAAH Reverse | GTA GCC TTT GTA GTG TTC CAT CT |
| DAGL Forward | GCT CTT CGG CTT GGT CTA TAA |
| DAGL Reverse | GCC ATT TCG GCA ATC ATA CAG |
| MAGL Forward | AAG AGT GGA GCG AGC AAT |
| MAGL Reverse | GAT GAT TCC ATG AGC AGG TAG G |

The primers were designed and bought in IDTdna.
